# ACUTE AMYLOID-Β 40 EXPOSURE DISRUPTS METABOLIC INSULIN SIGNALING IN BLOOD-BRAIN BARRIER ENDOTHELIAL CELLS

**DOI:** 10.1101/2025.11.21.689761

**Authors:** Douglas Nelson, Krishna R. Kalari, Karunya K. Kandimalla

## Abstract

**Background:** Brain insulin resistance and cerebrovascular dysfunction emerge early in late-onset Alzheimer’s disease, but how amyloid-β (Aβ) disrupts insulin signaling at the cerebrovascular blood–brain barrier—the principal site of insulin receptor signaling and transport into the brain—remains unclear.

**Methods:** We exposed two distinct human blood-brain-barrier endothelial cell models to soluble Aβ40 or Aβ42 for 1 h, followed by 100 nM insulin for 10 min. Protein and phosphoprotein responses were quantified by reverse-phase protein array, and differential expression was evaluated using empirical-Bayes-stabilized linear models with FDR correction.

**Results:** Aβ40 reduced insulin-stimulated Akt phosphorylation and converted insulin’s normal inhibition of AMPK into modest stimulation, increasing AMPKα Thr172 and ACC Ser79 phosphorylation. Aβ42 did not alter insulin-stimulated Akt signaling, indicating isoform-specific disruption of endothelial insulin responses. Without insulin, both Aβ40 and Aβ42 showed modest stimulation of ribosomal protein s6.

**Conclusions:** These findings show that Aβ40 acutely impairs insulin signal transduction in BBB endothelial cells, supporting a model in which vascular amyloid exposure contributes directly to the early development of brain insulin resistance in AD.

## INTRODUCTION

Alzheimer’s disease (AD), the primary cause of dementia in the elderly, is a multifaceted neurodegenerative disorder characterized by the accumulation of toxic amyloid-β (Aβ) peptides and neurofibrillary tangles of hyperphosphorylated tau [1–4]. In late-onset AD (after age 65) [5], these Aβ and tau pathologies consistently co-occur with cerebrovascular dysfunction and brain insulin resistance, both of which emerge early in the disease process [6–8].

Among the cerebrovascular changes associated with late-onset AD, dysfunction of the blood–brain barrier (BBB) is highly significant. The BBB forms the primary interface between blood and brain, regulating nutrient delivery and facilitating the clearance of metabolic waste, including toxic Aβ peptides [9]. The BBB is an integral component of the neurovascular unit (NVU), an interdependent association of neurons, astrocytes, pericytes, and the cerebral endothelium that couples neuronal activity to local changes in cerebral blood flow [10]. In Alzheimer’s disease, the BBB and broader NVU exhibit early and progressive dysfunction [11].

Metabolic syndrome and type 2 diabetes, which are characterized by peripheral insulin resistance, are major risk factors for AD [12]. Chronic peripheral insulin resistance disrupts canonical metabolic insulin signaling, leading to impaired energy metabolism and elevated inflammation that accelerate AD progression [13]. Notably, insulin receptors are enriched in brain microvascular endothelial cells compared to other NVU components, positioning the BBB endothelium as the dominant entry point and a major signaling portal for insulin action within the NVU [14]. Impaired insulin signaling at the BBB, as observed in Alzheimer’s models and human studies, is thought to disrupt neurovascular coupling and significantly reduce Aβ clearance from the brain [10,13,15]. These findings point toward a detrimental positive feedback loop in which Aβ impairs endothelial insulin signaling, leading to BBB insulin resistance, which in turn further limits Aβ clearance [13,15]. Therefore, there is an urgent need to elucidate precise molecular mechanisms underlying the role of insulin resistance and Aβ exposure in engendering BBB dysfunction.

The most abundant of the Aβ isoforms that accumulate in the AD brain, Aβ40, which predominates vascular amyloid deposits associated with cerebral amyloid angiopathy. In contrast, Aβ42 is the dominant component of parenchymal amyloid plaques [16]. Although less abundant overall, Aβ42 aggregates readily to form highly neurotoxic oligomers [17]. These two isoforms have been shown to exert distinct effects on the cerebrovascular endothelium [18]. Prior studies have demonstrated that Aβ interferes with insulin signaling in BBB endothelial cells and neurons; however, the precise underlying mechanisms and isoform specificity remain poorly understood in the BBB endothelium [15,19].

To dissect the direct effects of Aβ exposure, we examined the acute response to Aβ40 and Aβ42 on BBB insulin signaling. Identifying these transient molecular changes is useful because early signaling disturbances often initiate longer-term adaptive and degenerative processes, even though the resulting chronic phenotype may diverge from the acute response due to compensatory feedback or other complex biological responses [20,21]. Accordingly, these early signaling changes can provide insight into the initial conditions from which chronic endothelial dysfunction and brain insulin resistance can emerge. To capture these acute responses within a broader signaling context, we utilized reverse-phase protein arrays (RPPA). This antibody-based proteomic analysis enables the parallel measurement of hundreds of proteins and phosphoproteins, allowing us to interpret insulin pathway perturbations alongside changes in other vascular, metabolic, and stress-response nodes.

Gaps in understanding how Aβ disrupts BBB[22] insulin signaling remain a major obstacle to interpreting the characteristic vascular and metabolic pathophysiology of late-onset AD. Thus, identifying the early endothelial responses to Aβ exposure is crucial for understanding how vascular and metabolic dysfunction converge in AD and for advancing therapeutic strategies. Accordingly, this study systematically examines the acute, isoform-specific effects of Aβ40 and Aβ42 on BBB endothelial insulin signaling to characterize the direct molecular responses that may initiate these complex disease interactions.

## METHODS

### Aβ and insulin Peptides

Aβ40 and Aβ42 were obtained from Aapptec (Louisville, KY). Soluble Aβ40 and Aβ42 solutions were prepared according to the procedure described by Klien and characterized as described in our earlier publications [23]. Briefly, Aβ films were formed after dissolving the lyophilized peptide in HFIP and evaporating the solvent under vacuum. The resultant films were hydrated by sequential addition of 50 µL DMSO, 50 µL deionized water, and 100 µL pH 8.3 F12 medium, with brief sonication after each step. Then the solutions were centrifuged at 10,000 rpm for 5 min at 4 °C to remove any insoluble aggregates, and the supernatant was diluted in cell culture medium to final concentration of 1 µM, as assayed by ELISA. Vehicle controls were prepared using the same DMSO, water, and medium sequence without Aβ peptide.

Insulin powder (Sigma, Saint Louis, MO) was solubilized in 0.1 N sodium carbonate (5.3 mg/mL). A 1000 µM master stock was generated at 5.808 mg/mL and stored at 4 °C. From this, a final concentration of 100 nM was prepared in the cell culture medium, and the cells were treated with 100 nM insulin for 10 min.

### Cell cultures

Human cerebral microvascular endothelial cell (hCMEC/D3) line was a generous gift from P-O Couraud of the Institut Cochin (Paris, France). The endothelial cells were cultured according to the protocols provided by the Couraud group [22]. Polarized hCMEC/D3 cell monolayer were cultured by seeding the cells at passage 34 on 24 mm Transwells as detailed in our earlier publications [23].

Brain-specific microvascular endothelial cells (iBMECs) were differentiated from human induced pluripotent stem cells (hiPSC), according to Stebbins et al. and Motallebnejad et al [24,25]. The IMR90-4 line is derived from IMR90 normal human fetal lung fibroblasts and was reprogrammed by lentiviral delivery of OCT4, SOX2, NANOG, and LIN28. This female human iPSC line is distributed by WiCell as part of the IMR90 iPSC series. On day 8, iBMECs were transferred to collagen IV/fibronectin (CN IV-FN)–coated plates for selective endothelial attachment. After 1–2 h, unattached cells were removed, cells were washed once with Dulbecco’s phosphate-buffered saline, and the adherent endothelial cells were subcultured at 1 × 106 cells/cm2 onto 24 mm CellTreat® inserts (CELLTREAT Scientific Products, Pepperell, MA) and cultured for 1 day in endothelial cell medium to allow barrier formation. The endothelial medium consisted of Human Endothelial Serum-Free Medium (Thermo Fisher Scientific) supplemented with 1% fetal bovine serum (Thermo Fisher Scientific), 20 ng/mL basic fibroblast growth factor (PeproTech), and 10 µM retinoic acid (Millipore Sigma). After the first day, cells were maintained in Human Endothelial Serum-Free Medium containing only 1% fetal bovine serum. After 1 day, medium was changed to endothelial medium without retinoic acid and basic fibroblast growth factor.

For coculturing, frozen human brain vascular pericytes (hBVPs; ScienCell, Carlsbad, CA; passage 1) were thawed and expanded in gelatin-coated T-75 flasks using Pericyte Medium (ScienCell). Human astrocytes (HA; ScienCell, catalog #1800) were expanded per vendor instructions in Astrocyte Medium Complete Kit (ScienCell, catalog #1801). Pericytes and astrocytes were then seeded at a 1:1 ratio by cell number onto the bottoms of gelatin-coated 6-well plates (pericytes at 5,000 cells/cm2). After iBMECs formed a barrier, the inserts were placed into the pericyte/astrocyte plates to establish a non-contact BBB model; cultures were maintained at 37 °C in a humidified 5% CO2 atmosphere.

### Treatments with Aβ and insulin

Treatments and lysate preparation for iBMECs were the same as for hCMEC/D3 cells. At the end of each treatment, the spent medium was removed, and cells were washed twice with DPBS. Then DMEM (2.5 mL) was added to the abluminal side. One milliliter of either control medium or Aβ (1 µM) was added luminally and incubated for 1 h. Where indicated, cells were spiked with 100 nM insulin for the last 10 min before the termination of the experiment. All Aβ and insulin incubations were carried out on the luminal side.

Treatment groups included untreated medium, vehicle, insulin (10 min), Aβ40 (1 h), Aβ42 (1 h), Aβ40 (1 h) plus insulin, and Aβ42 (1 h) plus insulin. Each of the seven conditions had three replicates, totaling 21 wells across seven treatments. Plates 1 (untreated) and 2 (insulin) were processed in tandem; followed by Plates 3 (Aβ40 alone) and 4 (Aβ40 plus insulin); Plates 5 (Aβ42 alone) and 6 (Aβ42 plus insulin); and plate 7 (vehicle only). These groupings defined four independent RPPA runs (batches).

At the end of treatment, plates were placed on ice. Cells were immediately washed once with ice-cold HBSS, then ice-cold HBSS containing protease and phosphatase inhibitors was added to all wells. Then RIPA lysis buffer containing protease and phosphatase inhibitors (50 µL per well) was added. Cells were scraped with a cell scraper and collected in pre-chilled microcentrifuge tubes. This sequence was repeated for all wells. Lysates were sonicated for 4 min until clear and centrifuged at 10,000 rpm for 10 min at 4 °C. Supernatants were transferred to fresh 0.5 mL tubes, pellets discarded, and samples stored at −20 °C. For final preparation, 4X SDS without bromophenol blue and with beta-mercaptoethanol was added, samples were boiled for 5 min, and stored at −80 °C.

### Reverse Phase Protein Array (RPPA) assays

RPPA was performed by the MD Anderson Functional Proteomics RPPA Core (Set151; 21 samples; 306 antibodies). Lysates were blotted as five two-fold dilutions (undiluted, 1:2, 1:4, 1:8, 1:16) on nitrocellulose slides, probed using tyramide amplification, developed with DAB, scanned on a Huron TissueScope, and quantified in Array-Pro. Relative protein levels for each antibody were estimated by SuperCurve interpolation of the five-point dilution series. The Core provided sample-specific loading correction factors CF1 and CF2, where CF1 is calculated within the submitted set and CF2 across all samples printed on the slide; samples with CF2 < 0.25 or > 2.5 are flagged as protein-loading outliers recommended for exclusion. The hCMEC/D3 samples had CF1 0.83–1.35 and CF2 0.45–0.72, whereas iBMEC samples had CF1 0.66–1.33 and CF2 0.56–1.10. None of the samples met CF2 outlier criterion and were not excluded.

### RPPA data analysis

Sample-wise median centering was applied after inspection of raw expression distributions and variance profiles across samples. Probes corresponding to proteins that are not biologically relevant to the blood–brain barrier endothelium were excluded. The exclusion list was curated from established literature and cross-referenced for lack of functional abundance based on median values from pseudobulk analysis of a public transcriptomic dataset (NCBI GEO: GSE163577). A list of filtered probes is provided in Supplementary List 1. A total of 268 probes were included after filtering, out of which 62 were phospho-probes. Three filtered probes were relevant to pericyte biology and were not removed from the pericyte/astrocyte analysis.

Principal component analysis (PCA) was performed for each two-group contrast to assess replicate clustering before differential expression analysis. Cluster coherence was quantified using silhouette widths computed on the principal components explaining ≥95% of the variance, and Mahalanobis distances in two dimensions using a pooled within-class covariance estimate.

To reduce the influence of high-variance and outlier probes, differential expression was fit with Limma using empirical-Bayes moderation configured with a mean–variance trend (trend=TRUE) to accommodate RPPA’s intensity-dependent variance and robust hyperparameter estimation (robust=TRUE). We estimated sample precision weights with arrayWeights for each affected contrast.. Benjamini-Hochberg multiple hypothesis correction was applied per contrast.

To generate composite scores that emphasize probes supported by both meaningful effect size and statistical evidence, we transformed differential expression statistics into bounded scores using raised-cosine functions with log -fold change (LFC) parameters log2(1.1) and log2(1.5), and FDR parameters -log10(0.10) and -log10(0.001). Composite scores were then calculated as the geometric mean, penalizing imbalance between the two components.

To evaluate inter-batch shifts, we defined a set of invariant total-protein probes based on within-batch comparisons. A probe was considered invariant if, within each of the three two-group batches (untreated vs insulin, Aβ40 vs Aβ40 + insulin, Aβ42 vs Aβ42 + insulin), the estimated log fold-change between groups had a 95% confidence interval entirely within ±0.1. Confidence intervals were calculated from the limma model using empirical Bayes–moderated standard errors. Only probes meeting this criterion in all three batches were retained. For each inter-batch group comparison, we summarized the distribution of differences across the invariant set by reporting the Wilcoxon signed-rank p-value, the median log fold-change, and the median absolute deviation (MAD).

## RESULTS

### Differential expression overview and quality control

We evaluated replicate coherence and treatment separation prior to differential expression using principal component analysis. hCMEC/D3 contrasts involving Aβ plus insulin showed reduced coherence and greater within-group variability (Supplementary Table 1), so weighted linear modeling was applied to these contrasts. As a high-level summary of treatment effects, differentially expressed probes (FDR < 0.05) were tallied by direction across the eight contrasts for both endothelial models (Table 1).

**Table 1:** Counts of differentially expressed probes (FDR < 0.05) by direction (up = positive LFC; dn = negative LFC) in hCMEC/D3 (HD3) and iBMEC (IPS-derived) models across eight key contrasts.

| Contrast | HD3up | HD3dn | HD3tot | IPSup | IPSdn | IPStot |
| --- | --- | --- | --- | --- | --- | --- |
| vehicle-untreated | 27 | 76 | 103 | 2 | 7 | 9 |
| Aβ40-vehicle | 59 | 5 | 64 | 38 | 27 | 65 |
| Aβ42-vehicle | 34 | 9 | 43 | 7 | 9 | 16 |
| insulin-untreated | 30 | 52 | 82 | 14 | 28 | 42 |
| Aβ40 insulin shift | 7 | 10 | 17 | 45 | 25 | 70 |
| Aβ40ins-insulin | 5 | 13 | 18 | 20 | 12 | 32 |
| Aβ42 insulin shift | 21 | 17 | 38 | 5 | 0 | 5 |
| Aβ42ins-insulin | 9 | 20 | 29 | 5 | 2 | 7 |

To assess the extent of non-correctable batch effects, we used a set of validated total-protein probes that were stable within each batch and examined their stability across batches (**Figure 1**). Probes were considered stable if their 95% confidence interval for the log_2_ fold-change between treatment groups in each of the three two-group batches fell within ±0.1. Probes with any plausible mechanism for acute Aβ sensitivity were excluded. Using these criteria, 12 invariant probes were identified in the hCMEC/D3 cells and 53 in the iBMECs. Consistent directional shifts in these invariant probes suggest batch effects strong enough to produce reproducible bias in one direction, even after median centering.

**Figure 1:**
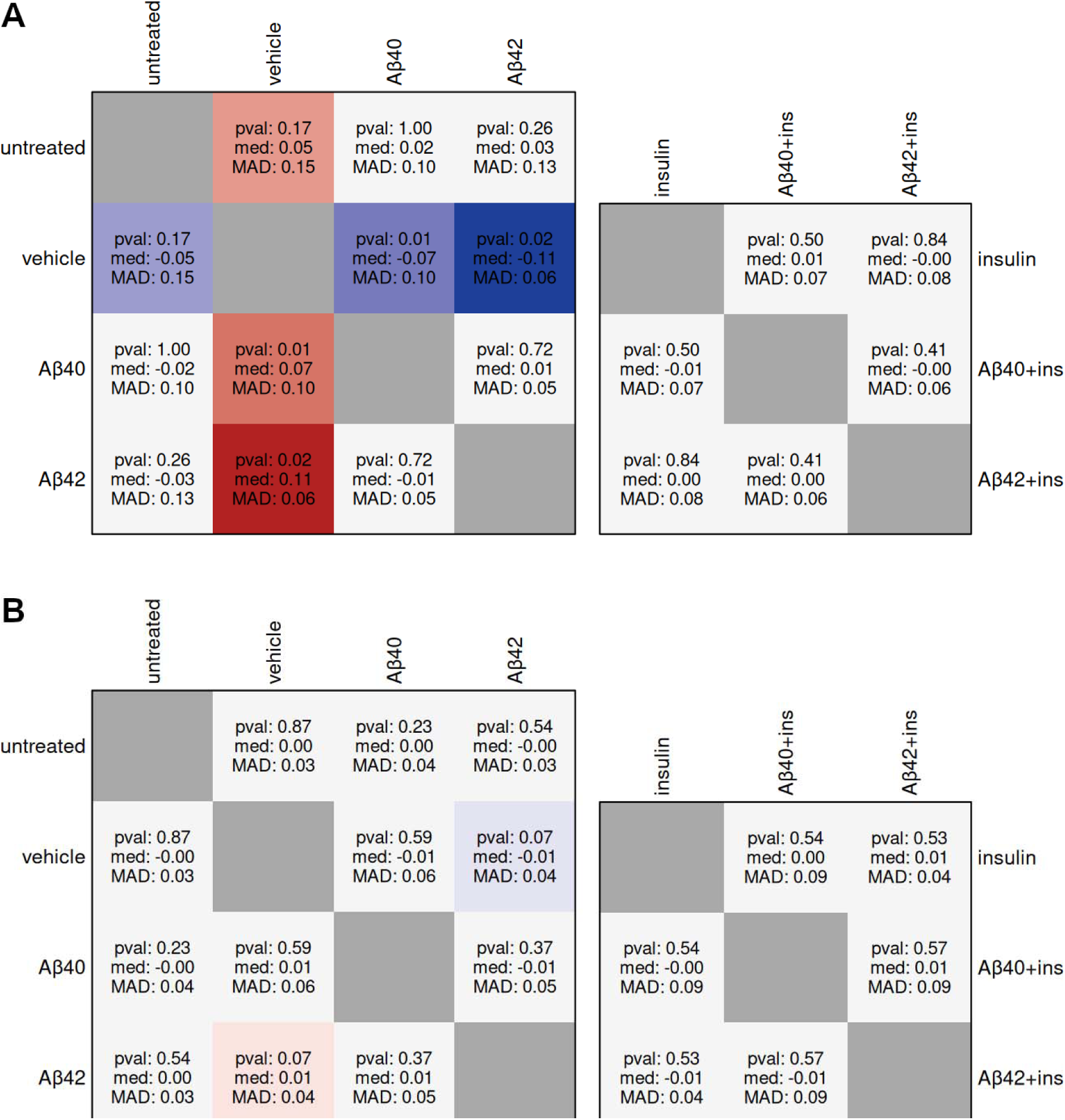
Evaluation of inter-batch shifts using invariant total-protein probes. RPPA analyses were run in four batches: (i) untreated vs insulin, (ii) Aβ40 vs Aβ40 with insulin, (iii) Aβ42 vs Aβ42 with insulin, and (iv) vehicle alone. An invariant probe set was defined as total-protein probes whose 95% confidence interval for log_2_ fold-change between treatment groups fell within ±0.1 in all three two-group batches, and this set was used to assess batch-to-batch consistency. Panel A (top) shows hCMEC/D3 results: the left matrix is a 4×4 comparison of all pairwise inter-batch shifts among the non-insulin groups (Untreated, Aβ40, Aβ42, Vehicle), and the right matrix is a 3×3 comparison among the insulin-containing groups (Insulin, Aβ40 with Insulin, Aβ42 with Insulin). Panel B (bottom) shows the corresponding iBMEC results with the identical structure. Each cell reports three statistics (top-to-bottom): Wilcoxon signed-rank p-value (pval), median log_2_ fold-change (med), and median absolute deviation (MAD). Cells are colored only when pval < 0.20 and |med| ≥ 0.01. Red indicates positive shifts and blue indicates negative shifts, defined as the row group minus the column group; color intensity scales with the magnitude of the median. Diagonal cells represent comparisons of a batch to itself and are shaded gray.

In the hCMEC/D3 model, the vehicle group (batch 4) showed a clear negative shift relative to the other batches, along with increased variability. Batches 1–3 were mutually stable, especially batches 2 and 3. Untreated versus Aβ comparisons showed moderate variability (median absolute deviations of 0.10–0.13) but no directional shift. Insulin-related contrasts—insulin alone or combined with Aβ40 or Aβ42—were consistent across batches, as were Aβ40 versus Aβ42 comparisons. Overall, batch 4 introduced a systematic negative bias, making vehicle-based contrasts less reliable. In the iBMEC model, the larger invariant probe set reflected lower overall variance, and batch-to-batch comparisons were generally stable, with only a slight negative shift in batch 4 relative to batch 3 (Aβ42).

Venn overlap analysis of DE probes (**Figure 2**) is consistent with a negative batch shift in the hCMEC/D3 vehicle run. Panel A shows that the vehicle–untreated contrast yielded 103 DE probes in the hCMEC/D3 monolayers, versus 9 in iBMECs (hiPSC-EC). Panels B and C also show that in hCMEC/D3 cells, most DE probes from the Aβ–vehicle contrasts overlapped those from vehicle–untreated (Aβ40: 78%; Aβ42: 63%), and all but one reversed log fold-change sign. All these overlaps were negative in vehicle–untreated and positive in Aβ–vehicle, consistent with systematically lower values in the vehicle group. Panels D and E show that the two Aβ isoforms exhibited modest concordance across both endothelial models: more than half of Aβ42 DE probes were also DE under Aβ40.

**Figure 2:**
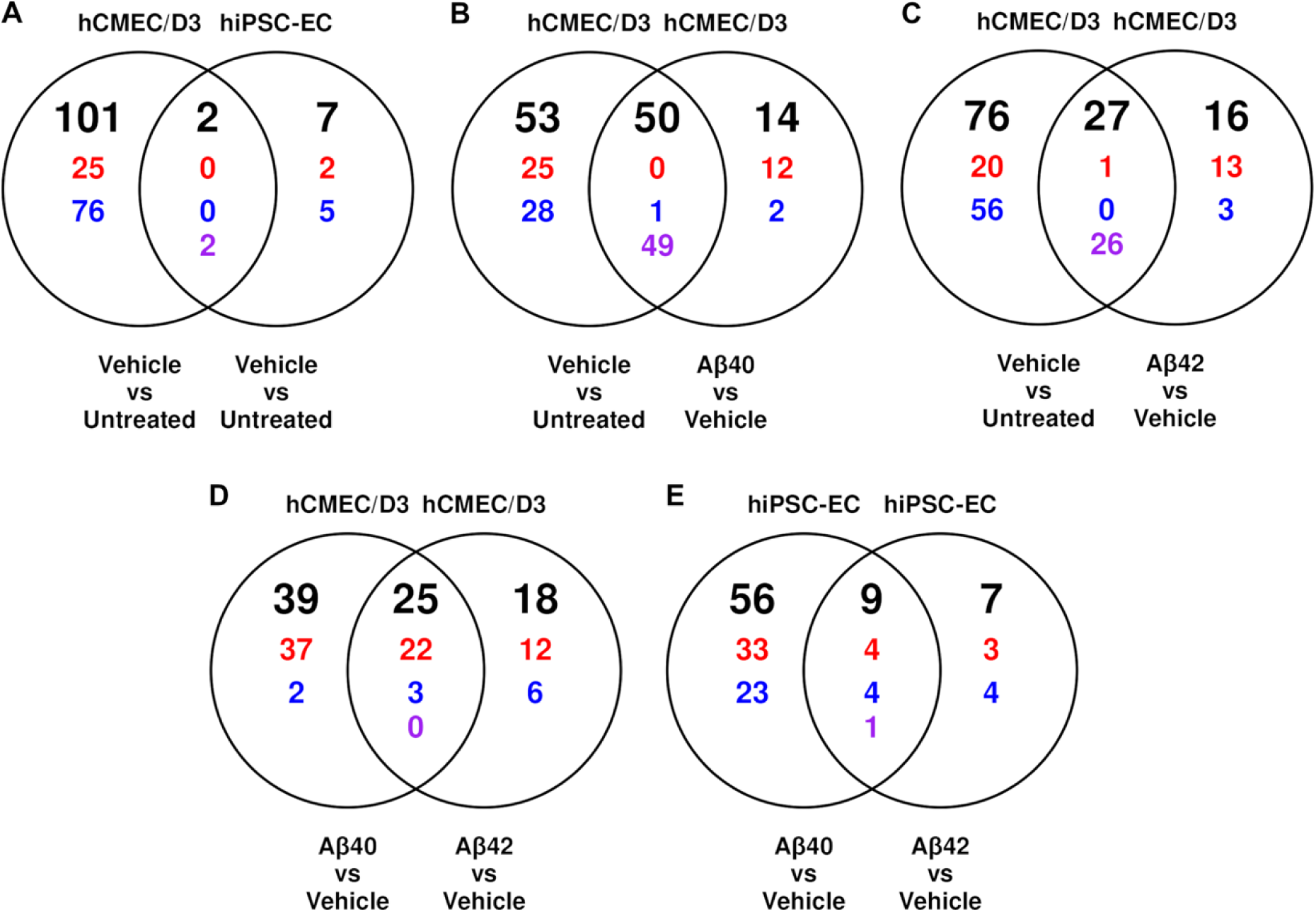
Venn diagrams of vehicle and amyloid-isoform effect contrasts. Differential expression is defined by Benjamini–Hochberg FDR < 0.05, with direction of change based on the sign of the log□ fold-change (first group minus second). Black numbers above each diagram indicate the total number of differentially expressed proteins. Colored numbers denote directional overlap: red for up/up, blue for down/down, and purple for opposite-sign overlaps. Left: DMSO vehicle vs untreated in hCMEC/D3 endothelial cells, compared with the same contrast in iBMECs (hiPSC-ECs) co-cultured with primary human pericytes and astrocytes. Middle: in hCMEC/D3, vehicle vs untreated compared with Aβ40 vs vehicle. Right: in hCMEC/D3, vehicle vs untreated compared with Aβ42 vs vehicle.

### Aβ effects without insulin

Figure 3 summarizes the top phosphoprotein responses to one-hour amyloid exposure in both BBB endothelial cell models, shown for both Aβ–vehicle and Aβ–untreated contrasts. The score matrix integrates fold-change and FDR into a single metric that reflects magnitude and statistical reliability. Ribosomal protein S6 (S6) Ser235/236 was upregulated in both contrasts and for both isoforms in hCMEC/D3 cells, while Ser240/244 was additionally upregulated in iBMECs with Aβ40. In iBMECs, AMPK Thr172 and ACC Ser79 were also upregulated by Aβ40, whereas AMPK Thr172, Akt Ser473, and GSK3α/β Ser21/9 were downregulated by Aβ42. In hCMEC/D3 cells, Aβ42 likewise downregulated Akt Ser473 and GSK3α/β Ser21/9, as well as Akt Thr308, although these changes were only observed in the Aβ–vehicle contrast.

**Figure 3:**
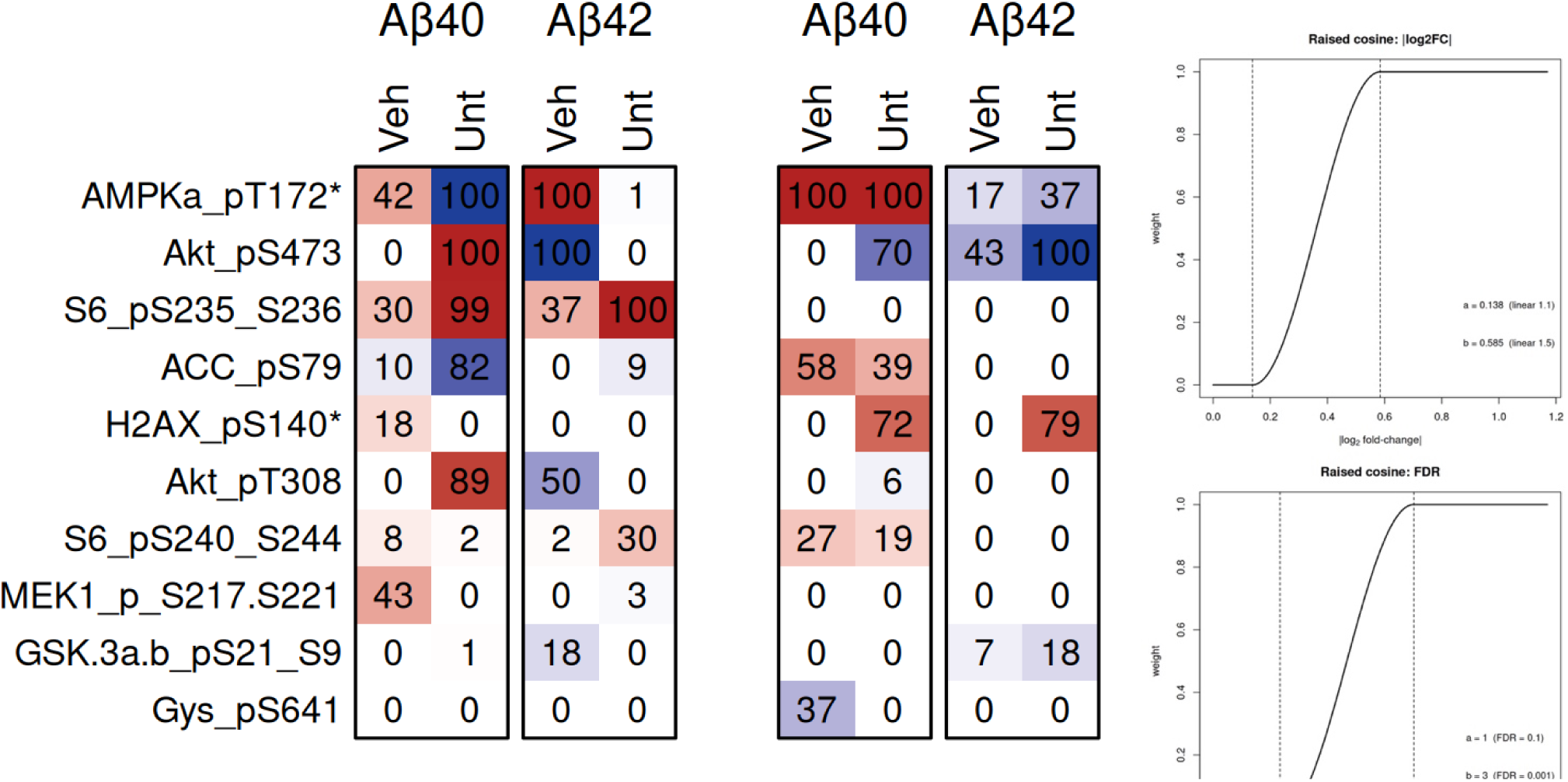
Differentially expressed phosphoprotein antibody probes in BBB endothelial models after one-hour Aβ exposure. Heatmap matrices show differentially expressed phosphoproteins in hCMEC/D3 (left) and iBMEC (right). For each Aβ isoform (Aβ40 and Aβ42), two contrasts are shown: Aβ vs Vehicle (Veh) and Aβ vs Untreated (Unt). Each cell contains a score computed as the geometric mean of two normalized functions—one based on log_2_ fold change and one on FDR—each scaled to [0, 1]. Scoring functions are shown at right. Red indicates upregulation and blue indicates downregulation. Only the ten highest-scoring probes are shown. Antibody probes marked with an asterisk (*) were not validated by the MD Anderson RPPA Core.

### Insulin signaling in hCMEC/D3 and iBMEC models

Figure 4 shows differential expression results after 10 min exposure to 100 nM insulin in the two BBB endothelial models. The displayed probes are grouped into four functional categories: metabolic (Akt pathway and immediate substrates), mTOR-centered (S6/S6K/TSC2/PRAS40/mTOR/Rictor), mitogenic (MEK–ERK cascade and adaptors), and AMPK. For each probe, we defined the expected direction of change at insulin stimulation from established literature; in the mitogenic set, both stability and increases were treated as consistent expectations. IGF1R_pY1135_Y1136 was included with an expectation of either increase or stability, given its relatively low insulin affinity. Phosphorylation of mTOR and Rictor is highly context dependent, so expectations for those sites were not assigned. Across probes, the observed log2-fold changes generally aligned with the normalized phospho-to-total protein signal.

**Figure 4:**
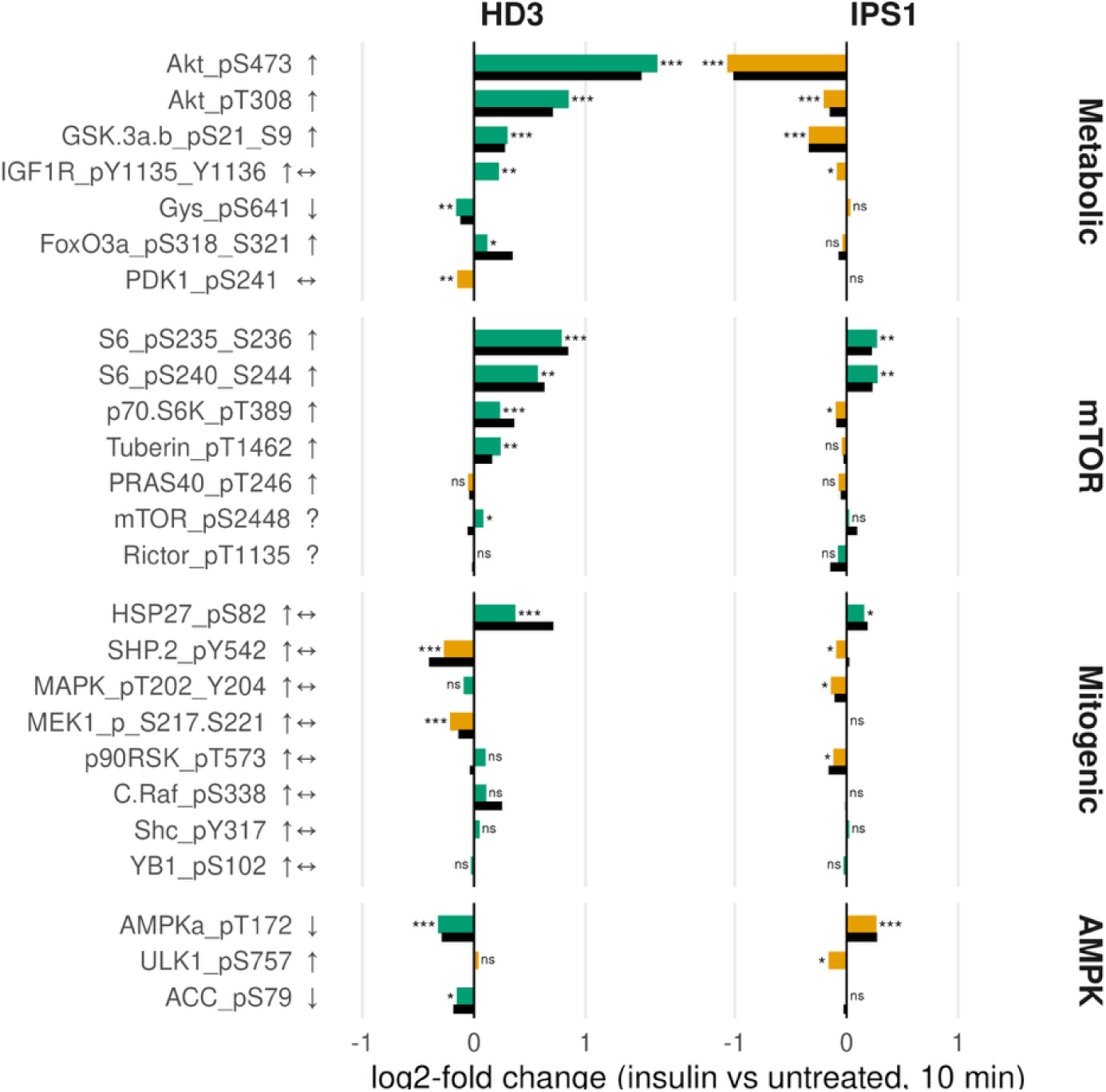
Log2-fold changes in insulin signaling probes in hCMEC/D3 and iBMEC BBB models. Cells were treated with 100 nM insulin for 10 minutes and compared with untreated controls. Probes are grouped into four functional pathways: metabolic, mTOR-centric, mitogenic, and AMPK. Expected direction of response is shown to the right of each probe (↑ increase, ↓ decrease, ↔ stable,no expectation). Colored bars represent observed log[-fold change: green = consistent with expectation, yellow/orange = divergence. *, **, *** denote FDR < 0.05, < 0.01, < 0.001; ns = not significant. Black bars represent the phospho-to-total normalized signal. Probes lacking a black bar lack a corresponding total-protein measurement (IGF1R_pY1135_Y1136, ULK1_pS757, Shc_pY317, YB1_pS102). For PDK1_pS241, the normalized value is near zero, consistent with its constitutive activation-loop phosphorylation.

The hCMEC/D3 model exhibits a canonical insulin response across metabolic and mTOR signaling nodes. Akt_pT308 and Akt_pS473 show the strongest activation, and downstream mTOR-axis substrates (S6_pS235/S236, p70-S6K_pT389, and Tuberin_pT1462) are similarly increased. In the AMPK group, AMPKα_pT172 and ACC_pS79 decrease as expected under insulin. In contrast, the iBMEC model shows attenuated and frequently discordant responses; for example, Akt_pT308 and GSK3α/β_pS21/S9 shift opposite to canonical expectations. Because only the hCMEC/D3 model demonstrates coherent pathway activation, all subsequent insulin-response analyses focus exclusively on the hCMEC/D3 system.

### Aβ effects on insulin signaling

In hCMEC/D3 cells, we used the second-order contrast (Aβ with insulin – Aβ) – (insulin – untreated) to quantify how Aβ modifies insulin-stimulated signaling (Figure 5). This contrast reports whether the insulin response at a given node is larger or smaller in the presence of Aβ. To determine whether these shifts represent augmentation, attenuation, or reversal of insulin action, we integrated the results of six additional contrasts (Table 2 and Table 3). This approach also reduces false-positive interpretation errors by identifying outcomes in which inconsistencies render interpretation inconclusive, or cases in which a reduced insulin response is equally compatible with attenuation or with saturation from an elevated baseline. The interpretive framework was formalized as decision trees, provided in Supplementary Figure 2. Results showed that Aβ40 produced the most substantial alterations in insulin signaling. Within the metabolic pathway, Aβ40 markedly blunted insulin-stimulated Akt activation at both pS473 and pT308 sites. In the AMPK group, Aβ40 reversed the expected insulin-mediated suppression of AMPKα_pT172, instead producing a net stimulatory effect. Both Aβ40 and Aβ42 produced modest reductions in insulin-stimulated mTOR-axis readouts (e.g., S6/S6K sites), although for Aβ40 this attenuation could reflect pathway saturation after Aβ40-alone stimulation.

**Figure 5:**
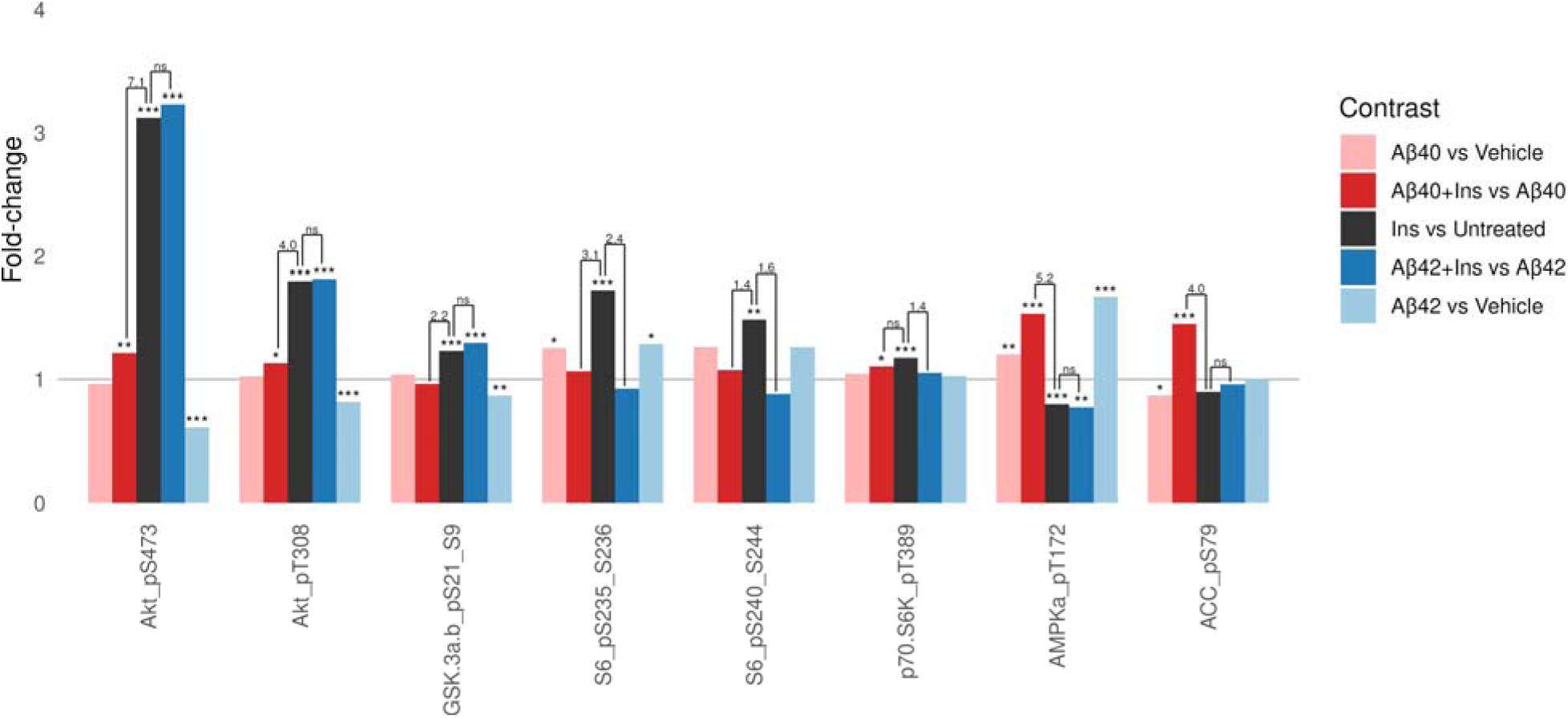
Insulin-responsive probes in hCMEC/D3 BBB endothelial cells with and without Aβ exposure. Shown are selected metabolic, mTOR-centric, and AMPK probes with significant insulin responses. Insulin effects are shown in untreated cells, and in cells exposed to Aβ40 or Aβ42 (both with DMSO vehicle). For reference, amyloid-vehicle contrasts without insulin are also shown. The horizontal line at fold-change = 1 denotes no change (log_2_ fold-change = 0) within each contrast. Asterisks (*, **, ***) denote FDR < 0.05, < 0.01, and < 0.001, respectively; ns = not significant. Numeric annotations above comparison brackets indicate -log_10_(FDR) from contrast-of-contrast tests, each bracket denoting the significance of differences between the effect of insulin on untreated cells and its effect on Aβ-treated cells.

**Table 2:** Summary effects of Aβ40 on insulin signaling in HD3 cells by integrating 7 contrasts. For each probe we report seven signed scores using the same method in Fig. 2: |LFC| and −logLL(FDR) are mapped to [0,1] combined by geometric mean, the sign of LFC reapplied, and scaled to ±100. Contrasts: V (vehicle − untreated), A (Aβ40 − vehicle), N (Aβ40 − untreated), I (insulin − untreated), IL (insulin with Aβ40 − Aβ40), L (level difference: insulin with Aβ40 − insulin), and S (shift in insulin response: IL − I). Interpretations are assigned by the decision rules in Supplementary Figure 2 (insulin response) and Supplementary Figure 3 (amyloid baseline effects).

| Probe | V | A | N | I | IΔ | L | S | Interpretation |
| --- | --- | --- | --- | --- | --- | --- | --- | --- |
| Akt_pS473 | 100 | 0 | 100 | 100 | 39 | -100 | -100 | Strong attenuation of insulin stimulation. |
| Akt_pT308 | 83 | 0 | 89 | 100 | 4 | -17 | -100 | Attenuation of insulin stimulation. |
| GSK.3a.b_pS21_S9 | 0 | 0 | 1 | 54 | 0 | -15 | -59 | Attenuation of insulin stimulation. |
| S6_pS235_S236 | 2 | 30 | 99 | 100 | 0 | 0 | -100 | Attenuation of insulin stimulation, or saturation following basal stimulation. |
| S6_pS240_S244 | 0 | 8 | 2 | 91 | 0 | 0 | -26 | Attenuation of insulin stimulation, or saturation following basal stimulation. |
| p70.S6K_pT389 | 0 | 0 | 0 | 33 | 1 | -1 | 0 | None. |
| AMPKa_pT172 | -100 | 43 | -100 | -61 | 100 | 37 | 100 | Reversal of insulin inhibition. |
| ACC_pS79 | -15 | -10 | -83 | -1 | 99 | 35 | 100 | Reversal of insulin inhibition. |

**Table 3:** Summary effects of Aβ42 on insulin signaling in HD3 cells by integrating 7 contrasts. For each probe we report seven signed scores using the same method in Fig. 2: |LFC| and −log_10_ (FDR) are mapped to [0,1] combined by geometric mean, the sign of LFC reapplied, and scaled to ±100. Contrasts: V (vehicle − untreated), A (Aβ40 − vehicle), N (Aβ42 − untreated), I (insulin − untreated), I[(Insulin with Aβ42 − Aβ42), L (level difference: insulin with Aβ42 − insulin), and S (shift in insulin response: I[− I). Interpretations are assigned by the decision rules in Supplementary Figure 2 (insulin response) and Supplementary Figure 3 (amyloid baseline effects).

| Probe | V | A | N | I | I□ | L | S | Interpretation |
| --- | --- | --- | --- | --- | --- | --- | --- | --- |
| Akt_pS473 | 100 | -100 | 0 | 100 | 100 | 0 | 0 | No effect on insulin response. |
| Akt_pT308 | 83 | -50 | 0 | 100 | 100 | 0 | 0 | No effect on insulin response. |
| GSK.3a.b_pS21_S9 | 0 | -18 | 0 | 54 | 73 | 0 | 0 | No effect on insulin response. |
| S6_pS235_S236 | 2 | 37 | 100 | 100 | 0 | -3 | -91 | Attenuation of insulin stimulation. |
| S6_pS240_S244 | 0 | 2 | 30 | 91 | 0 | -9 | -45 | Attenuation of insulin stimulation.<br>Basal stimulation by Aβ. |
| p70.S6K_pT389 | 0 | 0 | 0 | 33 | 0 | -28 | -2 | Weak inhibition of insulin response. |
| AMPKa_pT172 | -100 | 100 | -1 | -61 | -58 | -1 | 0 | No effect on insulin response. |
| ACC_pS79 | -15 | 0 | -9 | -1 | 0 | 0 | 0 | No effect on insulin response. |

## DISCUSSION

In this study, we examined how acute Aβ exposure affects insulin signal transduction in BBB endothelial cells. BBB insulin resistance in late-onset AD has frequently been associated with reduced insulin receptor abundance and inhibitory serine phosphorylation of IRS proteins. However, reductions in insulin receptor (INSR) abundance appear to develop gradually over the course of disease and have been linked to BACE1-dependent receptor degradation, rather than to immediate effects of Aβ exposure [26]. Consistent with this, total INSR protein levels were unchanged over the 1 h exposure interval. Although the acute signaling changes observed here are not consequences of receptor depletion, this does not rule out changes in receptor localization or availability. We therefore consider two mechanistic possibilities that are both consistent with our results and prior work: (1) direct interaction of Aβ40 with the insulin receptor, which could alter receptor availability or INSR signal output, and (2) inhibitory serine phosphorylation of IRS proteins driven by kinase pathways engaged during Aβ exposure.

### Direct Aβ-INSR interactions

Experiments using human placental membranes and purified insulin receptor preparations showed that Aβ40 and Aβ42 reduce insulin binding and inhibit insulin-stimulated receptor autophosphorylation in a manner consistent with competitive inhibition, at concentrations in the low micromolar range [27]. Competitive binding assays using purified insulin receptor preparations demonstrated that Aβ40 and Aβ42 can engage the insulin-binding ectodomain with substantially lower affinity than insulin, but within concentrations relevant to pathological Aβ accumulation [27]. Structural modeling of monomeric Aβ40 supports an energetically favorable but weaker interaction at the receptor ectodomain [28].

It is important to note that lower affinity, as in the case of IGF1 versus insulin, does not simply produce a weaker version of the insulin response [29]. INSR activation involves coordinated conformational changes and multisite autophosphorylation that determine the balance between downstream signaling branches; thus, different ligands can elicit qualitatively distinct signaling profiles even when binding at the same site. By the same logic, Aβ binding to INSR is unlikely to mimic insulin action at lower potency but may instead induce a distinct or incomplete signaling state whose downstream consequences remain poorly characterized.

Insulin binding to INSR—and, to a lesser extent, IGF1R or hybrid receptors—activates insulin receptor substrates (IRS), which recruit PI3K to the membrane. PI3K converts PIP2 to PIP3, enabling co-localization of PDK1 and Akt. PDK1 phosphorylates Akt at Thr308, and mTORC2 further activates Akt via Ser473 phosphorylation. Fully active Akt then phosphorylates and inhibits GSK3α/β at Ser21/9, an event that can also indirectly suppress AMPK through Akt-mediated phosphorylation of AMPKα at Ser485/491. This topology provides a defined framework for interpreting how Aβ alters insulin signal transduction in BBB endothelial cells.

The impact of Aβ on this cascade was examined using reverse-phase protein arrays (RPPA), which provide relative, semi-quantitative measures of protein and phosphoprotein abundance. Because RPPA platforms exhibit batch-dependent intensity offsets, the experiment was structured so that all insulin–versus–no-insulin contrasts were evaluated within the same array batch. To assess the reliability of comparisons across batches, we applied an invariant-probe analysis to estimate the residual directional displacement remaining after standard preprocessing.

### Assessing batch effects

The invariant probe set analysis (Figure 1) revealed a uniform negative displacement in the vehicle run, consistent with a remaining batch-specific directional bias. The observed negative displacement in the vehicle batch implies that differential expression results involving vehicle contrasts which are consistent with this direction should be interpreted with caution. For example, the higher number of differentially expressed proteins in hCMEC/D3 cells compared to iBMECs for contrasts involving vehicle samples are an expected result of this batch shift. However, several trends involving vehicle contrasts are most likely not confounded by this batch bias and therefore likely represent genuine biological effects.

The stronger Aβ40 than Aβ42 response cannot be attributed to batch effects, since the pattern appears in both cell models and, within hCMEC/D3, both contrasts share the same vehicle reference (Figure 1). Second, phosphorylation of Akt at S473 and T308 is robustly increased in the vehicle–untreated comparison, which is opposite in direction to the vehicle batch shift (Table 1 and Table 2). The first pattern aligns with the greater vasculotropic character of Aβ40 relative to Aβ42, while the second is consistent with reports of DMSO-induced Akt activation [31,32] In addition, basal levels of Akt at S473 and T308 were downregulated in Aβ42 compared to vehicle (opposite the vehicle batch shift), reversing DMSO-induced stimulation mentioned previously.

### Basal (not insulin-stimulated) Aβ effects

Figure 3 evaluates Aβ40 and Aβ42 treatments against both vehicle and untreated groups to separate Aβ-associated changes from vehicle-linked batch effects. Because the hCMEC/D3 vehicle run exhibited a negative shift, increases observed only in the Aβ–vehicle contrast (and absent in Aβ–untreated) are confounded by the batch shift. Conversely, changes present only in the Aβ–untreated contrast that moved in the same direction as the vehicle–untreated comparison are consistent with vehicle (DMSO) effects. Only changes showing the same directional shift in both Aβ–vehicle and Aβ–untreated comparisons were therefore interpreted as Aβ-driven.

Comparison of the hCMEC/D3 (HD3) and iBMEC (IPS) matrices in Figure 3 also indicates little concordance between the two BBB endothelial models: across hCMEC/D3 (48 differentially expressed probes) and iBMECs (49 probes), only 16 overlapped (∼17%), and half of these were directionally discordant.

In the hCMEC/D3 model, ribosomal protein S6 phosphorylation increased with both Aβ40 and Aβ42 in the Aβ–vehicle and Aβ–untreated contrasts, with stronger effects at Ser235/236 than at Ser240/244 (Figure 3). Because S6 is a direct target and canonical readout of the mTORC1–p70S6K pathway, these changes are consistent with Aβ-induced mTORC1 activity [32]. The absence of detectable phosphorylation of p70S6K at Thr389 may reflect a kinetic mismatch, as this site is transient and can decay more rapidly than S6. In addition, S6 can also be phosphorylated by RSK downstream of MAPK/ERK, consistent with the comparatively stronger Ser235/236 signal, which can occur independently of mTORC1 [33].

Figure 3 also shows dephosphorylation of ACC Ser79 with Aβ40 treatment, suggesting reduced AMPK activity, as ACC is a direct AMPK substrate [34]. Reduced AMPK activity would relieve inhibition of mTORC1 and would therefore be consistent with increased S6 phosphorylation [35]. Although decreased AMPK Thr172 phosphorylation is robust in the Aβ40–untreated contrast, the contradictory vehicle-reference result—consistent with a batch-specific offset—and the use of an antibody not validated for RPPA make this interpretation indeterminate. Differentially expressed total proteins were not included in

Figure 3 but are shown in Supplementary Figure 1. Given the brief 1-h exposure, transcriptional or translational regulation is unlikely to account for changes in total protein abundance. Although rapid degradation effects cannot be completely excluded, no affected target stands out as a likely candidate for such short-term regulation. Thus, these apparent differences most likely reflect batch variation, technical noise, or altered antibody accessibility.

### Insulin-induced signaling in hCMEC/D3 and iBMECs

Before assessing Aβ effects on insulin signaling, we examined the direct effect of insulin itself (100 nM, 10 min vs untreated) to confirm expected pathway activation. The 10-minute interval captures a signaling phase in which phosphorylation states change rapidly, yet is within the period when many metabolic signaling nodes—particularly those downstream and exhibiting sustained activation, such as Akt and S6—are expected to remain active [36]. In hCMEC/D3 cell monolayers, probes associated with PI3K–Akt, AMPK, and mTOR signaling follow a canonical insulin response pattern, with Akt phosphorylated at Ser473 and Thr308 showing the largest increase, consistent with amplified signal transduction through the PI3K–Akt–mTOR axis [37].

The decreases in AMPKα Thr172 and ACC Ser79 reflect a metabolic shift toward anabolism, while increases in S6 Ser235/236, S6 Ser240/244, p70S6K Thr389, and TSC2 (tuberin) Thr1462 are consistent with downstream mTORC1 activation [38,39]. IGF1R shows mild activation despite its low insulin affinity [40], consistent with the supraphysiological dose, while PDK1 Ser241 remains stable owing to its constitutive activation-loop phosphorylation [41]. Mitogenic (MAPK/ERK) pathway proteins show little or inconsistent change, likely reflecting the transient nature and context dependence of insulin-induced ERK activation, which is weak or absent in many endothelial and metabolic cell types [42–44]. Whereas hCMEC/D3 cells exhibit a canonical insulin response, the iBMEC model shows coherent opposite-direction changes across key metabolic nodes. This difference may stem from differences in growth-factor exposure between the two culture systems [45]. Hence, all subsequent analyses of Aβ effects on insulin signaling were limited to the hCMEC/D3 cell line.

### Aβ effects on insulin-induced signaling

To quantify how amyloid alters insulin responsiveness, we analyzed second-order intra-batch contrasts (Aβ with insulin – Aβ) – (insulin – untreated), which represent the change in insulin response after 1 h Aβ exposure (Figure 5). Final interpretation is made by integrating the results of these comparisons with other major contrasts (Table 2, Table 3 and Supplementary Figure 1). In hCMEC/D3, Aβ40 markedly attenuated Akt activation—most prominently at Ser473, and to a lesser extent at Thr308—whereas Aβ42 did not measurably alter Akt activation. Both isoforms appeared to blunt insulin-evoked mTOR-axis readouts (S6 Ser235/236 and Ser240/244, particularly the former), although these sites were already elevated at baseline after Aβ treatment, especially Ser235/236.

For S6 Ser235/236 and Ser240/244, the second-order contrasts show that the insulin-evoked increase was reduced in the presence of both Aβ isoforms. Aβ treatment elevated these phosphorylation sites at baseline, so the final levels with Aβ plus insulin were similar to those with insulin alone—and for Aβ40, statistically indistinguishable. This suggests the possibility of a ceiling (saturation) effect on S6 phosphorylation, rather than a genuine reduction in insulin responsiveness.

### Potential mechanisms of DMSO Akt stimulation

With respect to the observed DMSO-related Akt stimulation, our results are most consistent with an INSR-independent mechanism. Although DMSO has been shown to stimulate Akt phosphorylation in non-endothelial systems via a PI3K-dependent mechanism [32,56], the initiating transduction event remains unidentified. Several INSR-independent possibilities are plausible. DMSO perturbs membrane lipid order and raft organization, potentially activating PI3K/Akt through redistribution of raft-associated kinases such as Src, FAK, and integrins [57–59]. These considerations underscore the need to account for solvent effects and to evaluate alternative Aβ preparation methods, as both solvent and solute appear to exert confounding influences on the same key signaling node.

### Potential mechanisms of Aβ effects on insulin and AMPK signaling

Aβ exposure is widely reported to trigger oxidative stress, which can activate serine kinases such as JNK, IKKβ, PKCβII, and MAPKs that phosphorylate IRS at inhibitory sites [47–49]. RPPA results are inconclusive on this mechanism since the functional probes for IRS, IKK, and JNK2 were not part of the RPPA set, while JNK1 (Thr183/Tyr185), PKCβII (Ser660), and MAPK (Thr202/Tyr204) phosphorylations remained unchanged following 1 h Aβ treatment.

However, as noted before, Aβ-induced phosphorylation of S6 may reflect transient activation of S6K1 (p70S6K1) and/or MAPK–RSK during exposure that was not captured at the 1 h sampling point. Both of these can phosphorylate IRS at inhibitory sites, potentially independent of a ROS-induced stress response [50,51].

As discussed above, Aβ can engage the insulin-binding ectodomain of INSR. Aβ–INSR interactions have been shown to depend on aggregation state, with monomeric and oligomeric species exhibiting different receptor-binding behaviors and downstream signaling effects [29]. The distinct aggregation kinetics and pathways of Aβ40 and Aβ42 may underlie their divergent effects on insulin signaling. Aβ40 is relatively stable in monomeric form and, when it aggregates, proceeds more directly toward protofibrils and fibrils with little accumulation of soluble oligomers, whereas Aβ42 rapidly forms small, stable oligomeric assemblies [52]. We used a standard DMSO-based solubilization and dilution protocol known to yield predominantly monomeric Aβ40 and small soluble Aβ42 oligomers for short-term exposure assays [53–55]. Thus, aggregation-state differences offer a plausible explanation for the isoform-specific effects we observed—Aβ40 suppressing insulin-stimulated Akt activation, whereas Aβ42 does not and instead appears to neutralize the modest Akt stimulation produced by DMSO.

Along the AMPK branch, Aβ40 uniquely reversed insulin’s normal inhibition: phosphorylation of AMPKα Thr172 and its downstream target ACC Ser79 shifted from insulin-suppressed to insulin-stimulated. Under normal conditions, insulin restrains AMPK through Akt-mediated phosphorylation of AMPKα at Ser485/491, which prevents Thr172 activation. AMPK activation can occur through two principal routes: the canonical, energy-sensing (ADP/ATP ratio–dependent) LKB1 pathway and the Ca² /calmodulin-dependent CaMKK2 pathway.

## Conclusions

Our results showed that Aβ40 markedly attenuates insulin-stimulated Akt phosphorylation, thereby removing this inhibitory constraint and making AMPK more responsive to activating inputs. A likely source of such input is Aβ engagement of the RAGE receptor, which elevates reactive oxygen species and cytosolic Ca² to activate the CaMKK2–AMPK axis. CaMKK2 also stimulates eNOS, and the resulting nitric oxide transiently limits mitochondrial respiration, potentially adding an LKB1-dependent component to AMPK activation. While Aβ oligomers have been shown in several cell types to form Ca² -permeable pores that can rapidly increase cytosolic Ca², this mechanism is unlikely to contribute here, as monomeric Aβ40 is expected to be stable in the cell culture medium over 1 h.

In summary, Aβ40 appears to attenuate insulin activation of its main metabolic (Akt) signaling branch, while Aβ42 does not, suggesting direct INSR interaction with monomeric species as a potential factor. Additionally, our results indicate that Aβ40 may also uniquely reverse insulin’s canonical inhibitory effect on AMPK signaling, which could be due to the loss of Akt inhibitory phosphorylation of AMPK at Ser485/491 along with AMPK stimulation possibly mediated through an Aβ-RAGE- CaMKK2 route. These hypotheses are consistent with the observed activation of ACC immediately downstream of AKT. We have also shown that DMSO likely has confounding effects on Akt signaling, which raises the need to consider alternative solvents. And although the iBMEC model, which included pericyte-astrocyte co-culture, would generally be regarded as a more physiological model in several respects, its application in investigating insulin signaling needs to be carefully evaluated.

These findings show that short-term exposure to soluble Aβ alters insulin signaling in cerebrovascular endothelium, implicating direct amyloid effects in BBB endothelial insulin resistance. Of the two major isoforms, only Aβ40 disrupted endothelial insulin signaling, extending its known vasculotropic effects to include direct interference with metabolic signaling. This finding contributes to the emerging understanding that BBB dysfunction—and specifically endothelial insulin resistance—contributes to Alzheimer’s disease pathogenesis. Although the sample size was small and the RPPA platform inherently noisy, the principal results are qualitatively robust and provide a strong basis for future studies to define causal mechanisms—particularly receptor-level Aβ interactions, AMPK–Akt cross-regulation, and solvent effects.

## SUPPLEMENTARY

Supplementary List 1 : Removed probes unrelated to BBB endothelial biology: AR, Aurora.B, B7.H4, CD134, CD20, CD4, CD45, Claudin.7, c.Kit, E.Cadherin, EMA, ER, ER.a_pS118, GATA3, Granzyme.B, HER2, HER2_pY1248, HER3, HER3_pY1289, Heregulin, Lck, MS12, NAPSIN.A, Oct.4, P.Cadherin, PD.1, PR, Rab25, Syk, Tau, TTF1, UGT1A, UBAC1, ZAP.70, PLC.gamma2_pY759, MYH11, PDGFR.b, Notch3. Pericyte-related probes included in pericyte/astrocyte analysis: MYH11, PDGFR.b, Notch3.

---

**Supplementary Table 1:** Sample-level quality control metrics and the resulting Limma weights for the hCMEC/D3 contrasts identified as having reduced group coherence and increased within-group variability based on having Mahalanobis distances greater than 2.5 or negative Silhouette widths. The Limma weight applied to each sample is listed in the final column.

| Dataset | Contrast | Sample | Group | Mahalanobis dist. | Silhouette width | Weight |
| --- | --- | --- | --- | --- | --- | --- |
| HD3 | ab40ins_vs_ab40 | x7_3a_abeta_40_1hr | ab40 | 0.04695 | 0.19761 | 1.20 |
| HD3 | ab40ins_vs_ab40 | x8_3b_abeta_40_1hr | ab40 | 2.12002 | 0.28934 | 1.05 |
| HD3 | ab40ins_vs_ab40 | x9_3c_abeta_40_1hr | ab40 | 1.73742 | 0.14479 | 1.09 |
| HD3 | ab40ins_vs_ab40 | x10_4a_abeta_40_insulin | ab40_insulin | 2.63089 | -0.07614 | 0.33 |
| HD3 | ab40ins_vs_ab40 | x11_4b_abeta_40_insulin | ab40_insulin | 0.60798 | 0.16011 | 1.68 |
| HD3 | ab40ins_vs_ab40 | x12_4c_abeta_40_insulin | ab40_insulin | 0.85674 | 0.13219 | 1.31 |
| HD3 | ab42ins_vs_ab42 | x13_5a_abeta_42_1hr | ab42 | 0.46365 | 0.16072 | 1.02 |
| HD3 | ab42ins_vs_ab42 | x14_5b_abeta_42_1hr | ab42 | 0.00866 | 0.21246 | 2.09 |
| HD3 | ab42ins_vs_ab42 | x15_5c_abeta_42_1hr | ab42 | 0.36650 | 0.17958 | 1.11 |
| HD3 | ab42ins_vs_ab42 | x16_6a_abeta_42_insulin | ab42_insulin | 2.22765 | -0.06779 | 0.43 |
| HD3 | ab42ins_vs_ab42 | x17_6b_abeta_42_insulin | ab42_insulin | 2.66262 | 0.04448 | 1.15 |
| HD3 | ab42ins_vs_ab42 | x18_6c_abeta_42_insulin | ab42_insulin | 2.27092 | -0.00558 | 0.84 |

**Supplementary Figure 1:**
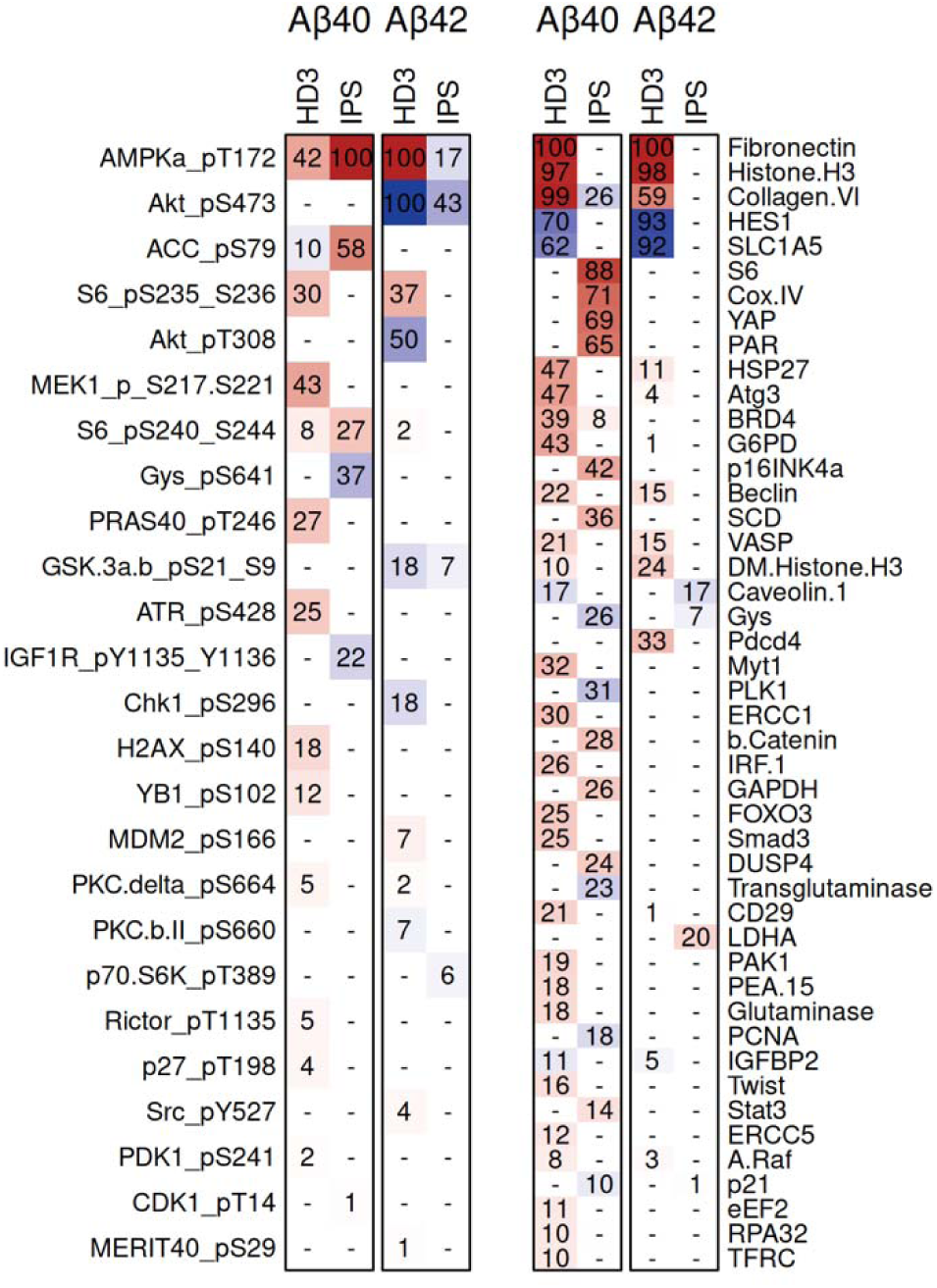
Differentially expressed probes in Aβ-vehicle contrasts with both isoforms and cell lines. Phospho-proteins are included in the right matrix and total proteins on the left.

**Supplementary Figure 2:**
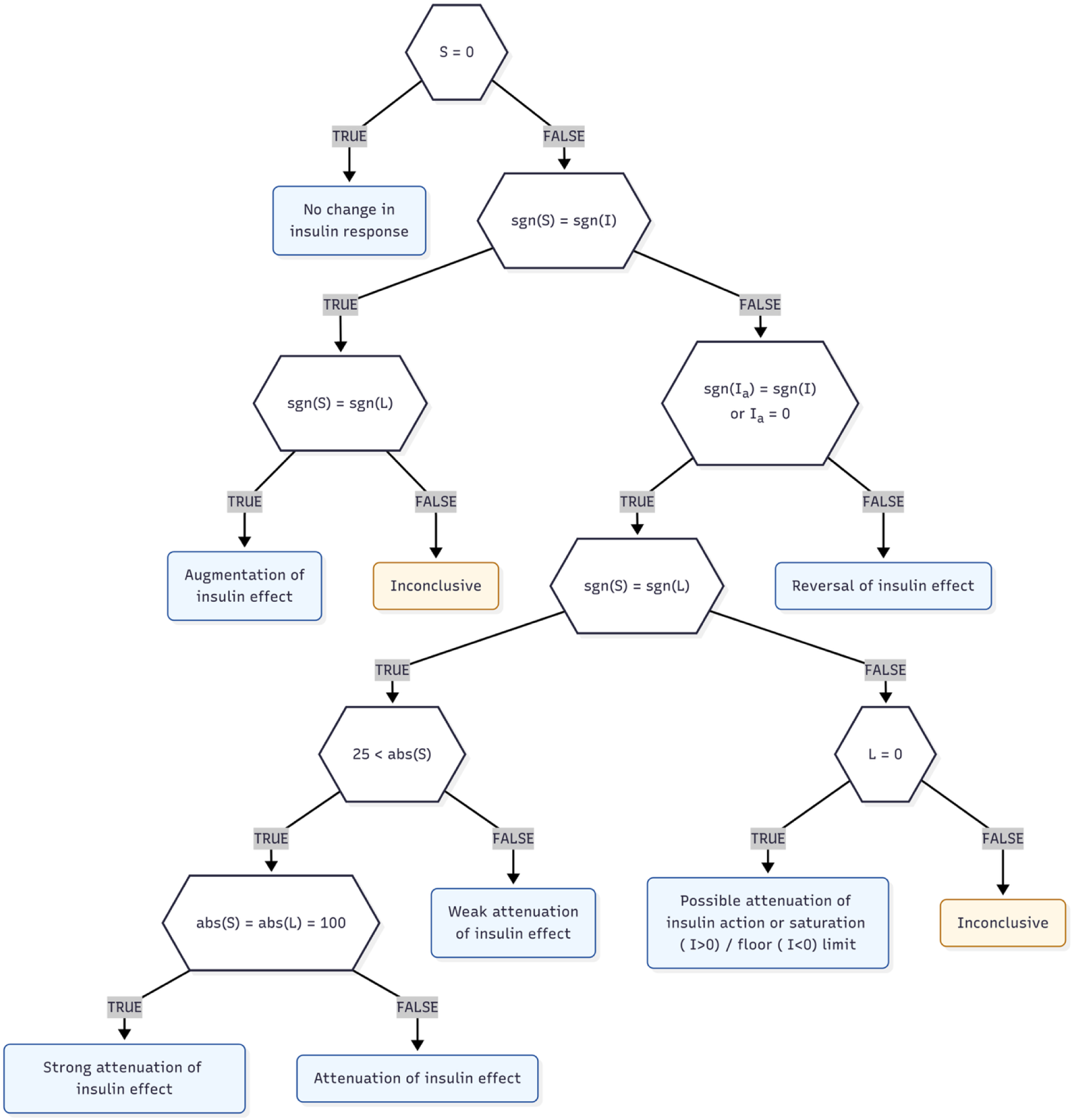
Decision tree for interpreting modulation of insulin response by Aβ using second-order and level contrasts. Contrasts are I = (insulin − untreated), I□ = (insulin with Aβ - Aβ), L = (Aβ+insulin - insulin), and S = I□ − I. Each contrast is discretized with the same scoring thresholds used in Figure 3, Table 2, and Table 3. Nonzero values require |log□ fold-change| > 0.138 and FDR < 0.1 (sign from LFC). The tree assigns: no change (S = 0); augmentation (sgn(S) = sgn(L)); reversal (sgn(I□) ≠ sgn(I)); or attenuation when sgn(S) = sgn(I) = sgn(L). Attenuation strength is qualified as weak if |S| < 25, strong if |S| = |L| = 100, otherwise unqualified. Cases with L = 0 under the attenuation branch are labeled “possible attenuation or saturation/floor.”

**Supplementary Figure 3:**
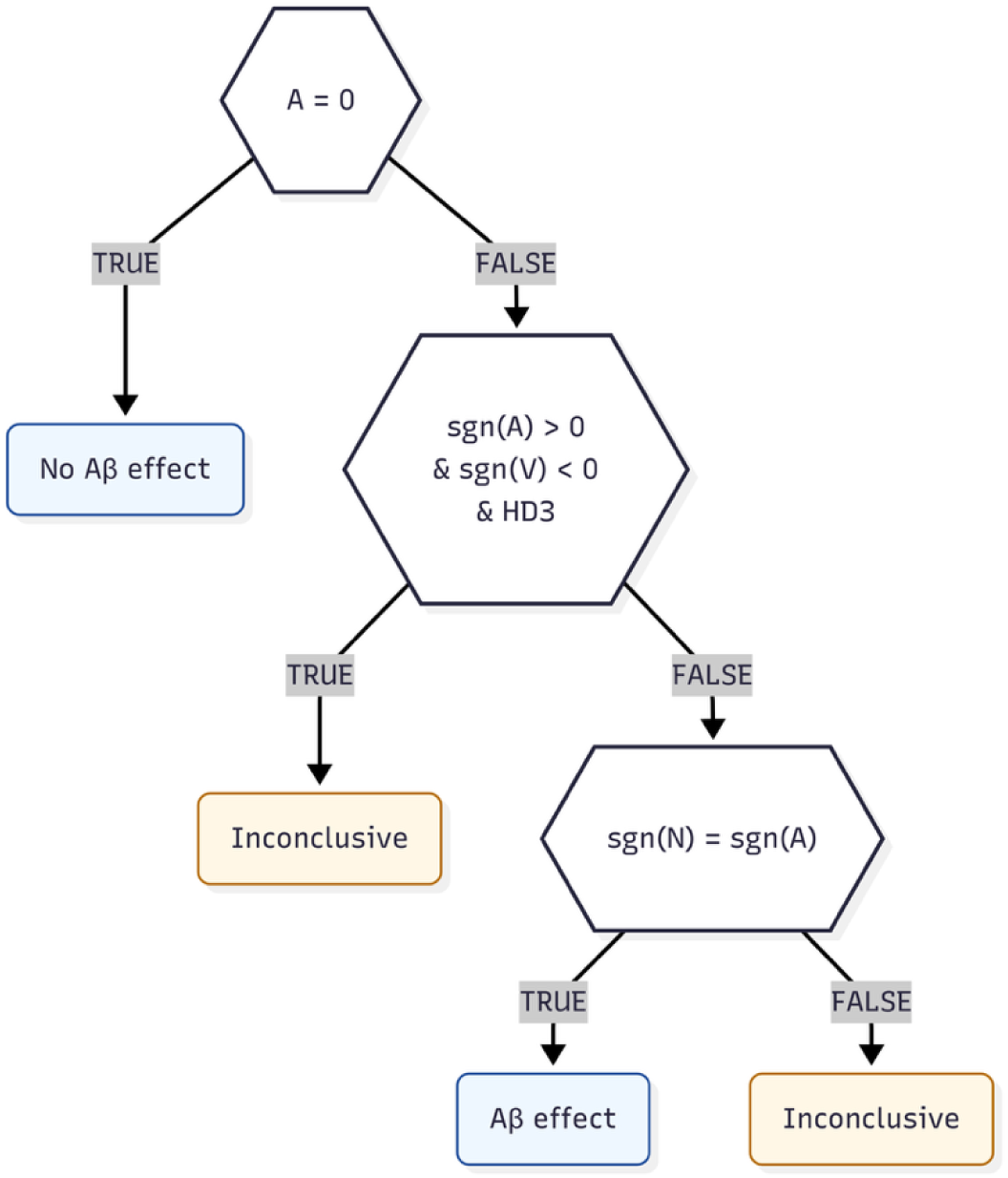
Decision tree for baseline Aβ effects in the absence of insulin. Contrasts are V = (vehicle - untreated), A = (Aβ - vehicle), and N = (Aβ - untreated), each discretized with the same thresholds used in **Figure 3**, Table 2, and Table 3 (|log□ fold-change| > 0.138 and FDR < 0.1 for nonzero values; sign from LFC). Aβ effect requires the change compared to vehicle to not be confounded by negative batch effects in HD3 vehicle and to have the same results in comparisons with untreated as to with vehicle.

**Supplementary Figure 4:**
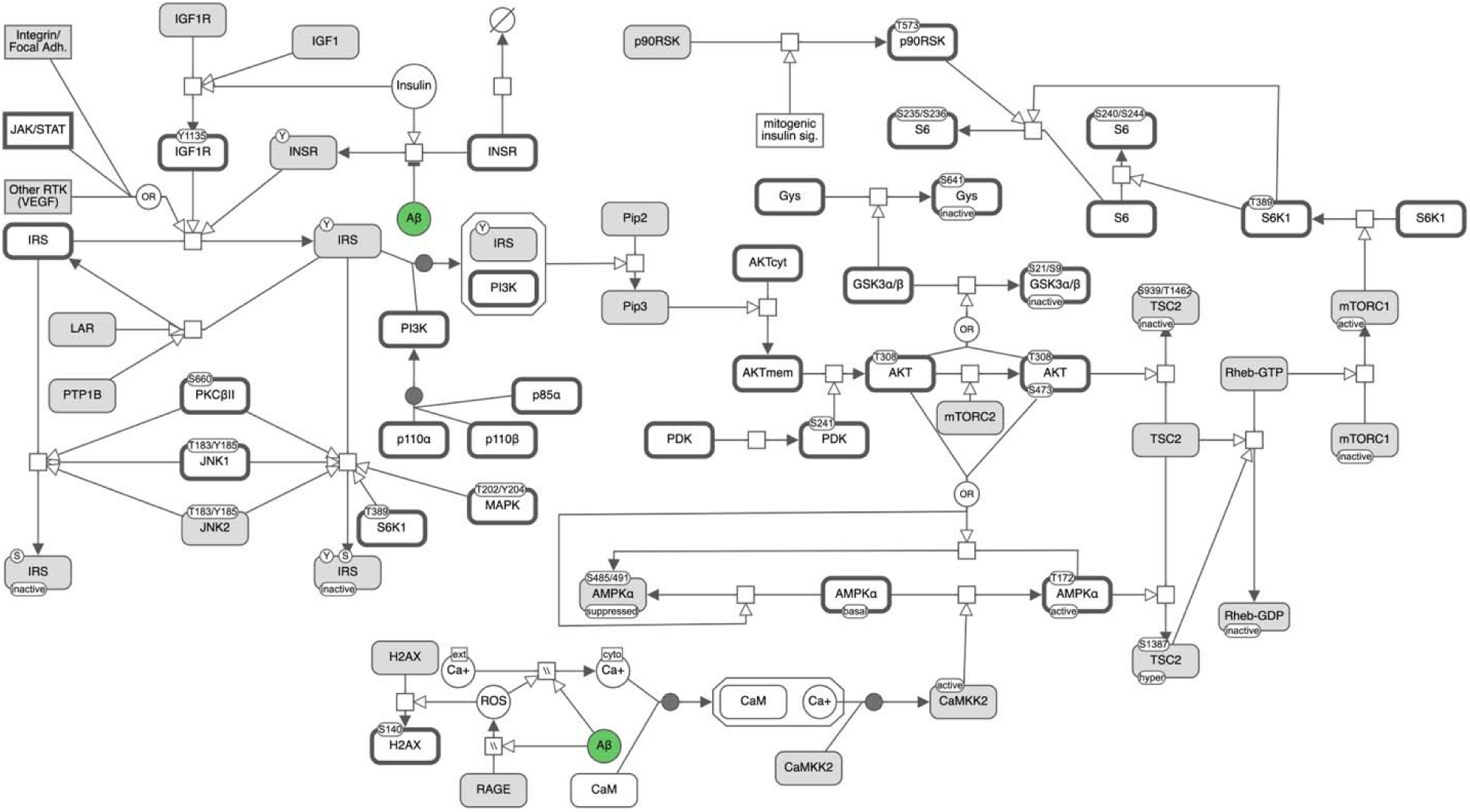
Systems Biology Graphical Notation (SBGN) Process Description graph of metabolic insulin signaling and related processes that includes measured RPPA probes in white and nodes without corresponding probes in grey.

